# Empowering future scientists: Mentors employ various strategies to engage students in professional science disciplinary literacy practices

**DOI:** 10.1101/2024.03.15.585231

**Authors:** Trisha Minocha, Tanya Bhagatwala, Gwendolyn Mirzoyan, Gary S McDowell, Sarah C Fankhauser

**Affiliations:** Emory University; Wake Forest University School of Medicine; Lightoller LLC, Ronin Institute, Institute for Globally Distributed Open Research and Education, School of the Art Institute of Chicago; Oxford College of Emory University

**Keywords:** Writing mentorship, disciplinary literacy, science writing

## Abstract

Peer-review and publication are important parts of the scientific enterprise, and research has shown that engaging students in such scholarly practices helps build their sense of belonging and scientific identity. Yet, these disciplinary literacy skills and professional practices are often part of the hidden curriculum of science research, thus excluding students and others from fully understanding ways in which scientific knowledge is constructed, refined, and disseminated even though students are participating in such activities. Secondary students are increasingly involved in scientific research projects that include authentic disciplinary literacy components such as research proposals, posters, videos, and scientific research papers. More and more, students are also engaging in professional practice of publishing their scientific research papers through dedicated secondary science journals. How teachers and other mentors support the development of professional disciplinary literacies in students is critical to understand as part of supporting more student participation in research. To this end, we used a mixed-methods study of interviews and surveys to examine the experience and conceptions of the mentors (teachers and professional scientists) who guided pre-college students through the writing and publication of their scientific research projects. Analyzing our data from a lens of cognitive apprenticeship, we find that mentors encourage independence by primarily employing the method of “exploration”. We also find that mentors have divergent views on the value of publication within science, versus for student scientists specifically. Our findings suggest that mentors could work to explicitly reveal their own thinking within science writing to provide more sequenced support for student scientists.

## Introduction

In the last two decades, research opportunities have become a common high school student experience. Science fairs, school-based and out-of-school research experiences, and summer science programs have increased significantly in the last ten years, with close to 1 million students participating in science fairs each year (Grinell, et al 2023). Pre-college research experiences can foster students’ research identity (Deemer et al., 2022); interest and creativity within STEM (MacLeish et al., 2012; Lin & Schunn, 2016 ); positive attitudes towards STEM (Vennix et al., 2018); and interest and retention in STEM through college (Dabney et al., 2012; Miller et al., 2018; Sahin, 2013; Kong et al., 2014; Maltese & Tai, 2011).

Such research opportunities often include *disciplinary literacy* outcomes, which emphasize developing skills gained from creating, communicating, and applying knowledge within a specific field (Shanahan & Shanahan, 2012). In scientific fields, this can involve various forms of communication practices such as writing research reports, developing proposals, emailing or dialoguing with key stakeholders, participating in peer review, crafting poster or oral presentations, or even participating in the broad dissemination of one’s work through publication (Fankhauser et al., 2021; Otto et al., 2023; Florence & Yore, 2004; Watts-Taffe et al., 2022). As emphasized by the Next Generation Research Experiences, science instruction and science research should include the disciplinary literacy skills of science, including how reading, writing, and thinking are part of the scientific process (Houseal et al., 2016).

Providing students the opportunity to participate in authentic disciplinary literacy practices allows them to practice employing the unique knowledge and skills that individuals use when engaging in the practices of their field (Shanahan & Shanahan, 2012). This has the potential to enhance literacy and identity factors. Participation in science literature can increase student understanding of science inquiry and concepts (Golden, 2023). At the undergraduate level, students who participate in authentic peer-reviews of preprint articles report greater STEM identity (Otto et al., 2023); and similarly at the graduate level, students who engage in the disciplinary literacy practices of writing a research paper and serving as a peer-reviewer report higher identity factors, indicating that students view participation in these practices as part of their scientific identity (Florence & Yore, 2004). Authentic engagement in the research process, as characterized by students proactively taking ownership of learning and demonstrating resilience in the face of adversity (Sinatra et al, 2015), may increase student confidence in scientific abilities which further strengthens their sense of identity in the STEM field (Trujillo & Tanner, 2017). When students are more engaged, the chances that they pursue a future in this discipline increases, especially with regards to underrepresented groups (Vincent-Ruz & Schunn, 2018; Chen et al., 2020). Our own research shows that pre-college students, including those who identify as underrepresented or disadvantaged in STEM, report gains in science identity, self-efficacy, confidence and belonging after participating in the publication of their research through the pre-college science journal: *[Journal]* (Authors 2021; Authors 2023; Authors 2022a).

Key to the successful outcomes of these student research experiences are the individuals who guide students throughout the research process, within or outside the classroom setting, including teachers, mentors, supervisors or sponsors.. Educators, peers, and families have an effect on student’s interests in STEM and this influence affects student’s beliefs on their own capabilities and career expectations (Nugent et al., 2015). Specifically, mentor support is essential to the deep learning associated with out-of-school research experiences such as science fairs (Andersson, et al., 2021; Behar-Horenstein et al., 2010). In one study, students reported that they felt “cared” for by dedicated research mentors, while mentors in the same study expressed the importance of their role in providing students access to resources and equipment (Andersson et al., 2021). Mentorship has been shown to be key to the development of underrepresented students, which has been associated with retention in the STEM field (Atkins et al., 2020). Underrepresented students in particular face distinctive barriers which can undermine their confidence and retention in STEM disciplines. Mentors play a crucial role in fostering support and guidance to navigate these challenges (Lisberg & Woods, 2018; Atkins et al., 2020; Ghazzawi et al., 2021). Successful mentors provide an environment where students are able to explore their interests as well as have a platform to succeed, positively influencing not only their academic careers but also their personal lives. In general, individuals are more likely to continue pursuing their academics when they are supported by someone they trust (Lisberg & Woods, 2018).

Research mentors need to guide students through activities *beyond* the experimental aspects of a research project. The professional disciplinary literacy practices of writing, peer-review, and publication are critical parts of the STEM research experience. Studies have shown that in tertiary education, students report “learning by doing” rather than specifically instruction on the theory, purpose, and strategies associated with professional disciplinary practices (Chong & Mason, 2021; McDowell et al., 2019; Gewin, 2019; van der Loo et al., 2019). Although students are able to learn by doing, processes and curricula that are hidden or more nuanced may require mentors to explicitly verbalize their thought processes and provide detailed explanations to support student understanding (Haeger & Fresquez, 2016; Jayabalan et al., 2021). As educators and researchers seek to make science practices more accessible, and as practices evolve and expand to include younger students, it becomes increasingly important to understand how to mentor students through different parts of the research process, especially disciplinary literacy practices.

Given that mentor perceptions and values can directly impact the belief and process development of the mentees, it is important to explore how mentors express these values (Nugent et al., 2015). For example, do mentors of younger students also take an approach of “learn by doing”, or are they more intentional about their approach to instructing and guiding students in disciplinary literacy practices?

Understanding how mentors mechanize and verbalize their values can provide insight for developing mentoring relationships. Although much research has been deployed to understand the mentor-mentee relationship and impact, no research presently exists (that we could find) on mentor perceptions and enactment of professional disciplinary literacy skills as part of the research process for pre-college students (García-Verdugo et al., 2020; Ian O’Byrne et al., 2021; Spires et al., 2018). To bridge this gap, it is essential that we explore the importance that mentors place on professional literacy practices, specifically for pre-college students, and how this guides them while mentoring students through the various aspects of the research process. In this study we seek to answer the following research questions:

● What are the mentors’ *perceptions* of the role of disciplinary literacy practices of peer-review and publication [Journal], particularly for the pre-college students they are mentoring?
● Why are mentors *motivated* to guide their students in [Journal] publication?
● How do mentors *support* their students in peer-review and [Journal] publication?
● What do the mentors *experience* in the mentorship process?

## Theoretical Framework

Our work emerges from the role of mentors in helping develop science identity, thus the theories we rely on are that of science identity and cognitive apprenticeship. According to Carlone and Johnson (2007), science identity has three critical components: recognition, competence, and performance.

Recognition occurs when both peers and the scientists themselves acknowledge their inclusion in the field. Competence refers to the depth of understanding of the subject matter. Performance is the ability to effectively communicate information according to established scientific standards (Carlone and Johnson, 2007).

Carlone and Johnson clearly illustrate that developing these components, and thus a strong science identity is closely linked to the persistence in scientific fields (Atkins et al., 2020; Chang et al., 2011; Chemers et al., 2011; Hazari et al., 2013; Perez et al., 2014; Woods et al., 2023).

Having a greater understanding of factors, activities, and experiences that contribute to the underpinnings of performance, competence and recognition, could give researchers and practitioners a roadmap for increasing persistence in STEM. For example, supportive mentorship is one way in which students may gain *recognition* for their scientific abilities. But what are the ways that students achieve external recognition? How might students develop competencies that are beyond experimental practices but still critical to being a successful researcher? And how can students engage in the vast array of norms of their discipline? Although Carlone and Johnson did not specifically include disciplinary literacy participation within their study or their model, we find that there is significant overlap between the outcomes of engaging in disciplinary literacy endeavors and the underpinnings of STEM identity (Figure 1). For example, students who participate in reading and writing of scientific articles are *performing* authentic skills of professional scientists. Students who submit a paper for peer-review, such as with the *Journal of Emerging Investigators,* increase their perceived *competence* in science writing and analysis (Fankhauser, et al 2021; Mattison, et al 2022). And importantly, through the peer-review and publication processes, students gain *recognition* by others in the science community for their scientific contributions. We and others have demonstrated that engaging in writing and publication of research papers facilitates growth in STEM identity, self-efficacy, and confidence, thus suggesting that enculturation in the disciplinary literacy norms of STEM could be drivers of STEM identity (Fankhauser, et al 2021; Mattison, et al 2022; Kim, et al 2023).

**Figure 1.**
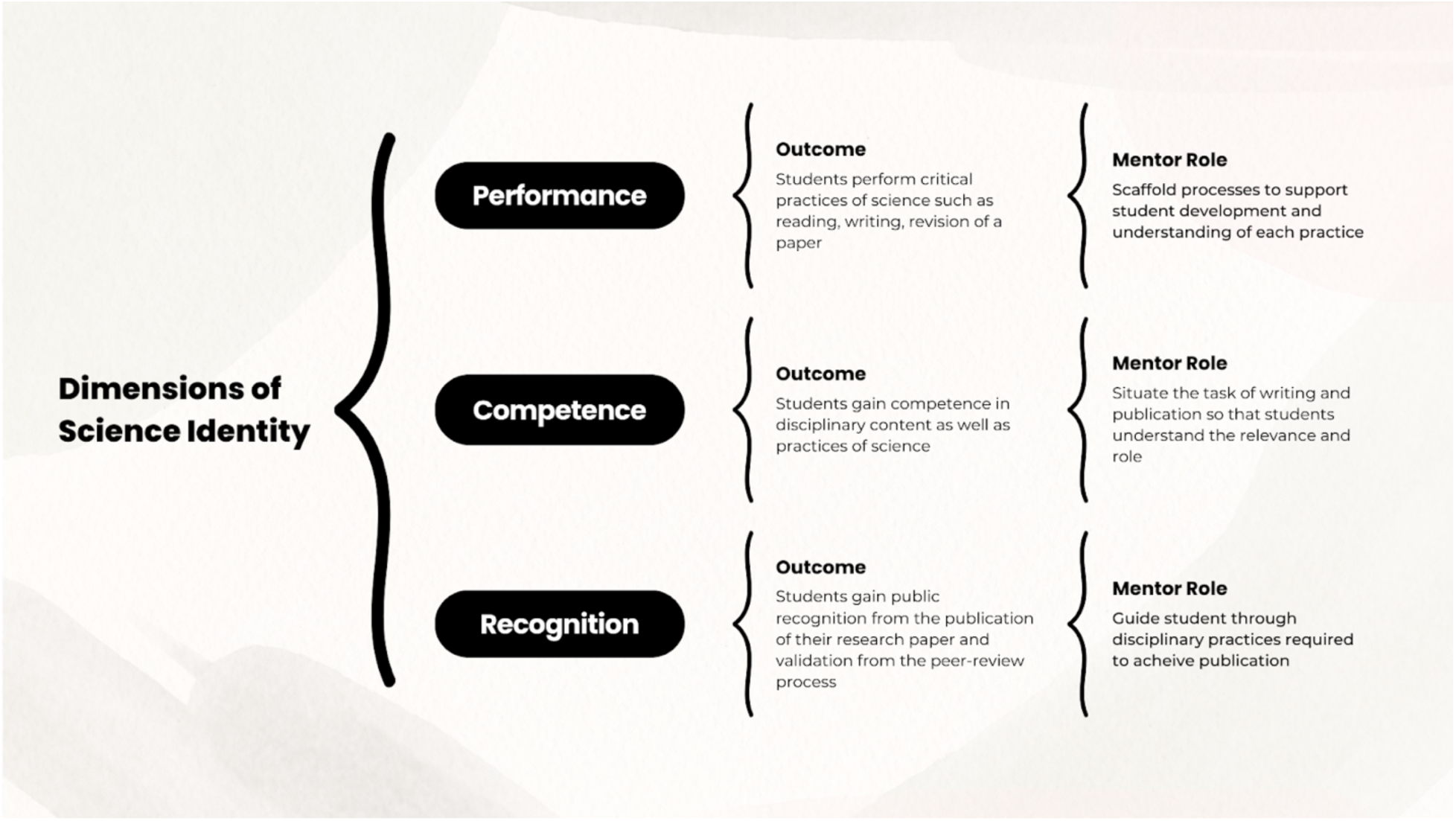
Three dimensions of scientific identity paired with the outcomes hypothesized to be achieved when students participate in the publication of their science research papers and the role of the mentor.

Like with other research practices, the role of mentoring can be critically important for the students’ development of competence in, and understanding of, disciplinary literacy practices within the larger scholarly enterprise (Ackerman, 1995; Keller et al., 2014; Casanave, 1992; Dysthe, 2002; Li, 2019; Kennedy, 2017). Considering the STEM identity theory put forth by Carlone and Johnson (2007), the role of a mentor expands to making the purpose and *performance* of disciplinary literacy practices visible such that the student becomes an insider to the community. This allows them to be successful in accomplishing the *practices* and gaining *recognition* (Hansen and Adams 2010). <u>As such, we draw on the construct of cognitive apprenticeship to explore the role of the mentor in developing students as enculturated members of science discourse.</u>

The theory of cognitive apprenticeship examines the role of a mentor in facilitating the entry of a newcomer into a disciplinary field, and the influence that quality mentorship can have on a student’s persistence in the field (Seymour, 2004). The four dimensions of cognitive apprenticeship: content, methods, sequencing, and sociology describe the types of knowledge, strategies to develop expertise, sequence of learning, and environmental contexts of learning, respectively (Collins, et al., 1991; Minshew, et al., 2021). These dimensions facilitate the key overarching aspect to cognitive apprenticeship: making the thinking of the discipline highly visible to the student/mentee. In other settings, cognitive apprenticeship has been a way to mentor students through a variety of academic and research processes (Stewart & Lagowski, 2003). These mentors have the role of demonstrating and coaching students through the activities while providing support and feedback to ensure students are able to effectively perform the skills learned themselves (Collins, et al., 1991; Minshew, et al., 2021). Not only has this pattern of coaching and scaffolding played a role in the enhancement of students’ academic skills, but it has also been successful with strengthening soft skills. A study that explored effects on students applying peer review skills learned in a classroom setting to a real-life situation, concluded that students greatly benefitted from peer reviewing an instructor’s literature review and showed increased levels of confidence, validating their knowledge of the learned peer review processes (Klucevsek, 2016). While students immersed themselves in the process of scientific communication by active participation, this type of learning refined their writing, critical thinking, and communication skills (Kolikant et al., 2006).

However, writing and peer review is not the end of professional practices: publication is an equally important process to scholarly endeavors. Publication can mean a variety of things depending on the field, but for this project we define publication as the process of submitting a paper for peer-review from an organized journal, subsequent revision based on reviewer feedback, and then dissemination on the journal’s platform whether that is online or in print. The chance to publish one’s research in a journal is generally viewed as important to one’s career for more advanced scholars, but it can also be a critical learning point for younger students who are beginning to navigate science and the science community (Fankhauser et al., 2021; Kim et al., 2023; Florence and Yore, 2004; Aitchison et al., 2010; Aitchison et al., 2012). Such experiences may facilitate or reinforce students’ confidence in the STEM field and accentuate their potential for future success (Paglis, Green & Bauer, 2006). Many report learning these processes by “doing”. However but without intentional approaches that meaningfully involve students in the steps of relevant disciplinary literacy practices, students are dependent on an advisor who “allows” shadowing or co-authoring, often leading to inequities in terms of involvement throughout the publication process (Abbott, et al 2020; Foster, et al 2012; Giuliano, et al 2022; Pratt, et al 2019). Cognitive apprenticeship offers an approach that allows students an opportunity to learn the mechanics and purpose of the process, while also experiencing the community involved in shaping scientific publications.

Although still limited, cognitive apprenticeship has been studied more at the graduate and postdoc level rather than the high school level (Stewart & Lagowski, 2003; McDowell et al., 2019; McDowell et al., 2021; Minshew et al., 2021; Blume-Kohout, 2017).

Our study focuses on the mentors who worked with pre-college students who were novices in the field of science publication. This posed unique challenges as novice researchers have not gained enough experience to independently manage the process of writing a paper. Basic writing skills, confidence, and organization of paper structure are some of the difficulties that novices face when learning to write in the discipline (Shah, et al 2009)., Learning to effectively engage in the discourse of the field and the norms of a disciplinary community could be facilitated when students participate in a cognitive apprenticeship at any experience level (Kolikant et al., 2006). When students are introduced to the specific strategies and disciplinary contexts of the field, they are able to be immersed in its practices and ways of thinking. By doing so, even young students may become familiar with genre conventions and communities associated with the genre. To be clear, we, the authors, do not equate students’ “success” with publication, but rather we value the process that students and their mentor go through, which serves to engage students in critical scholarly processes and may or may not produce a publication.

## Gap in field and Present study

While research has demonstrated both the positive impacts of mentoring and influence of participating in disciplinary literacy practices on STEM research students (Li, 2019), no study has investigated how mentors conceive of, and put to practice, the role of professional disciplinary literacy practices of peer-review and publication at the high school level. And although scientific writing mentorship has been studied at the undergraduate (Gamberi & Hall, 2019; Gruber et al., 2020; Reynolds & Thompson, 2011) and graduate levels (Florence & Yore, 2004; Thein & Beach, 2010; Aitchison et al., 2012), the ways in which mentors guide their students in writing and publication processes is not widely understood for younger students. Studies have shown that engaging in disciplinary literacy practices are key to helping students build their scientific identity, and mentors play a big role in students’ success in professional disciplinary literacy (Carlone & Johnson, 2007). By investigating mentors of pre-college students who published their research in the high school science journal, *[Journal],* our study aims to understand the motivation of mentors to guide students through the writing, peer-review, and publication processes, as well as their conceptions and mechanization of these processes. This will allow us to discover the role that mentors play for students engaging in disciplinary literacy. Finally, we look at what the mentors potentially gain or learn from guiding their students through these processes. By understanding these dimensions of mentoring, we hope to gain insight on ways to further strengthen mentoring of young scientists both in and out of the classroom.

## Methods

Our methodological framework followed a mixed-methods approach using survey responses and in-depth interviews of mentors (Yin, 2015). A survey allowed us to examine experiences and conceptions of publication from a larger population of mentors whose students had published with the [Journal]. A phenomenological analysis using in-depth interviews of a subset of mentors facilitated a deeper understanding of the survey responses and research questions (Charmaz, 2003).

### Acronyms

As described below we use the following acronyms throughout the paper: TPM: Teaching Profession Mentor SPM: Science Profession Mentor

### Participants and setting

[Journal] is an online science journal dedicated to peer-reviewing and publishing the research from middle and high school students. Students who submit papers are required to have an adult senior author, and this person may be a science teacher, university/college professor, or a guardian. Students come from a variety of demographic, geographic, and disciplinary backgrounds that have been explored in other studies (Fankhauser, et al., 2021; Mattison, et al., 2022; Kim, et al., 2023). We use the general term “mentor” to refer to the adult who helped the student author in their research and publication processes. For the interviews, mentors self-identified as being in a teaching profession or not, and we used this distinction to examine differences across the population. We use the subcategories of “teaching profession mentor” (TPM), to distinguish mentors who served as the student’s instructor within a school setting or described their role as education based. We use the term “Science Profession Mentor” (SPM) to refer to individuals who helped guide students in their research projects and did not identify as being in a teaching profession. All SPMs that we interviewed were in a scientific research profession. Respondent background and demographic data of survey respondents and interviews are shown in Tables 1 and 2, respectively.

**Table 1:** Demographic information for [Journal] mentors responding to survey

| **Demographic** | **Total** |
| --- | --- |
| **Race** |  |
| White | 40 |
| South Asian or Indian | 9 |
| East Asian | 7 |
| Southeast Asian | 1 |
| Black or African American | 2 |
| Latinx | 2 |
| Prefer not to answer | 3 |
| **Gender** |  |
| Female | 32 |
| Male | 31 |
| Prefer not to answer | 2 |
| **Highest Qualification Earned** |  |
| Bachelor's Degree | 5 |
| Some Graduate School | 4 |
| Masters Degree | 28 |
| Professional Degree | 28 |
| **STEM degree obtained** |  |
| Yes | 58 |
| No | 7 |
| **Prior participation in professional research** |  |
| Yes | 53 |
| No | 12 |
| **Prior experience in journal article publication** |  |
| Yes | 40 |
| No | 25 |
| **Prior experience in research mentorship** |  |
| Yes | 43 |
| No | 22 |
| **Educational Position** |  |
| Public/state school | 15 |
| Private school | 16 |
| International school | 1 |
| College/University Professor | 19 |
| Not employed in formal educational role | 14 |

**Table 2:**
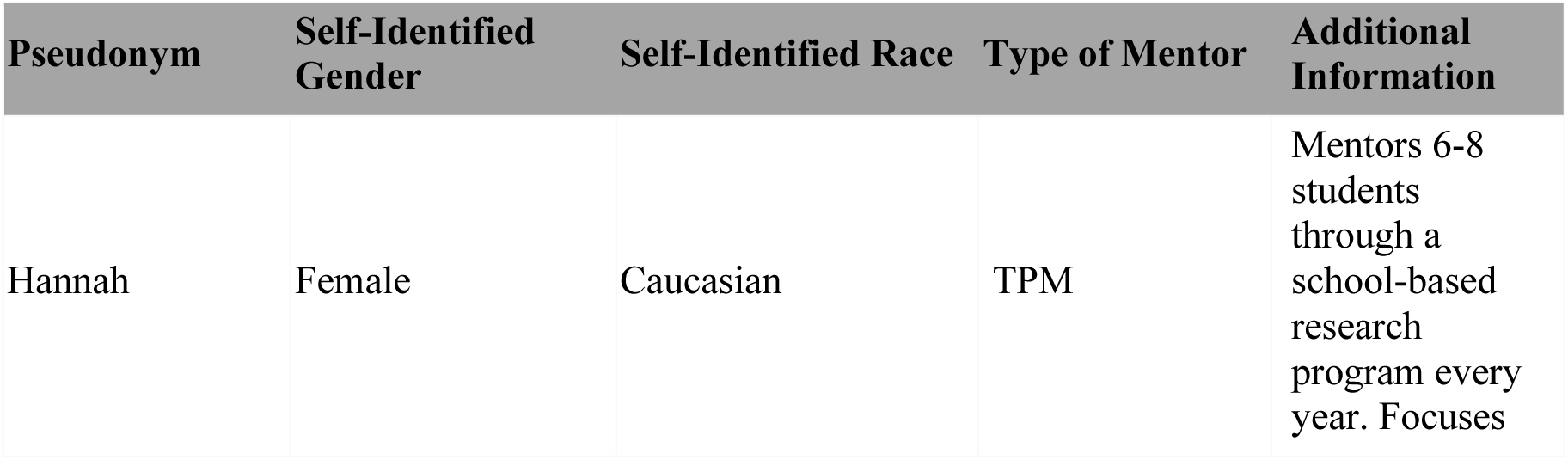
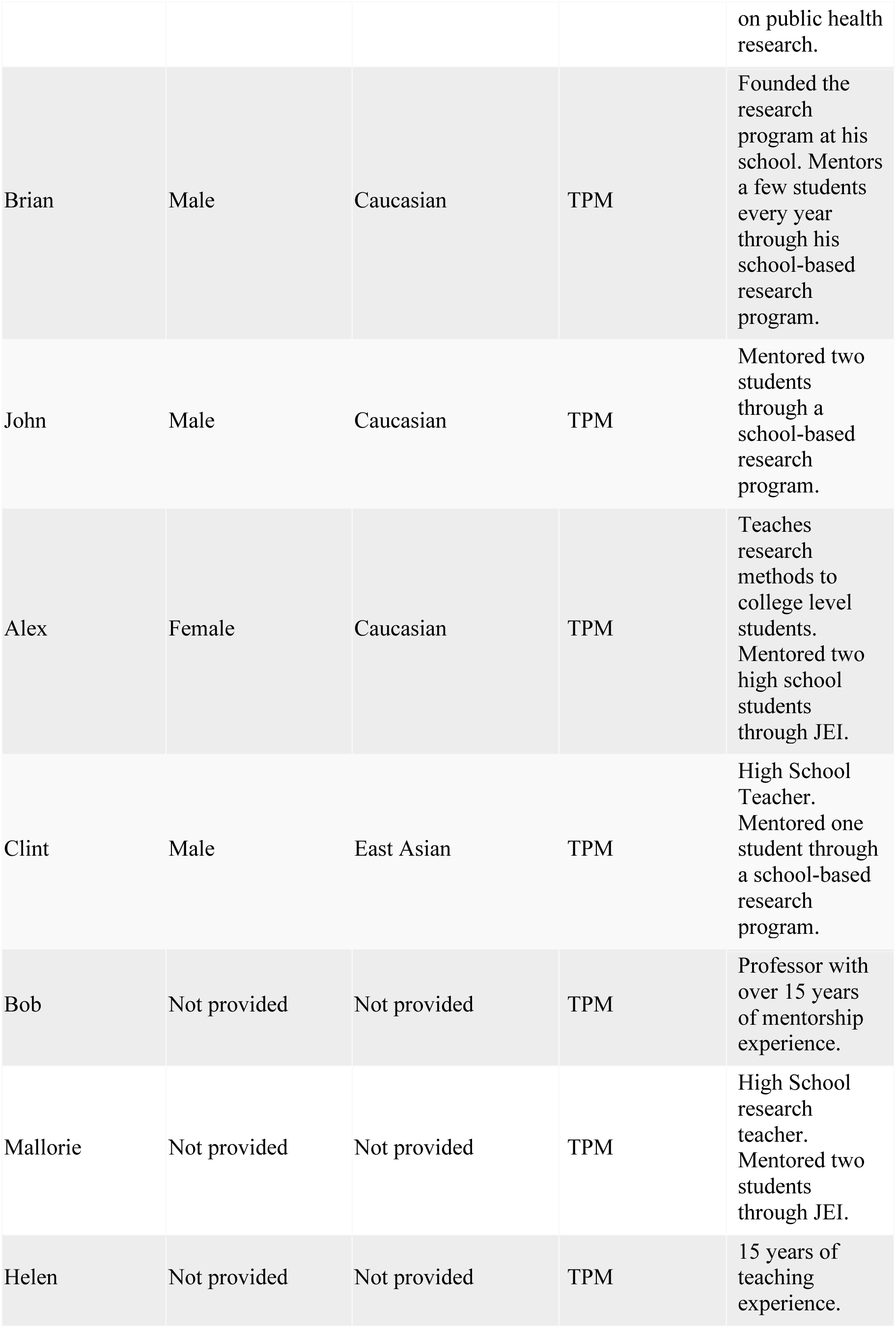
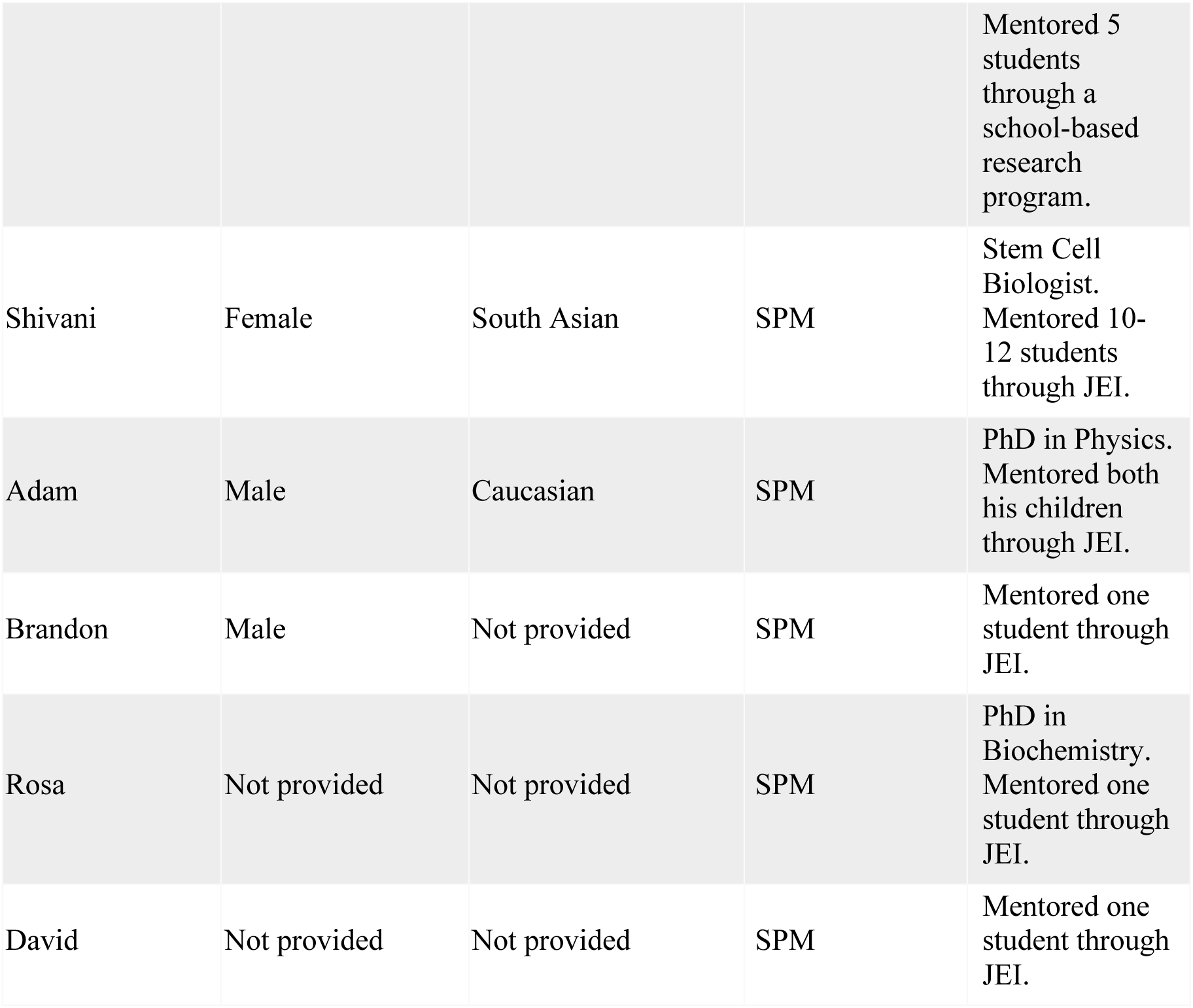
Demographic information for [Journal] mentors interviewed

### Survey Data collection

A survey was constructed in Qualtrics with a mix of Likert-scale and open-ended response questions asking the mentors about their previous experience with peer-review and publication, their confidence in these processes, and motivation and challenges associated with mentoring a student. Survey questions which assessed mentor science identity and self-efficacy were adapted from the Persistence in the Science Survey (Hanauer, et al., 2016). Additional questions were based on a survey previously used to assess perceptions of disciplinary literacy practices within science (Mattison, et al., 2022).Two identical versions of the survey were each deployed to different populations of student authors’ mentors. The first was sent to mentors who had students publish with [Journal] prior to 2020. Of the approximately 200 mentors emailed, 40 completed the survey. The other was sent to mentors as part of a final publication email, starting in late 2020. Upon receiving this email containing the final proof of the manuscript prior to online publication, mentors were asked to complete the voluntary survey. From 2020 to 2022 there were approximately 300 mentors contacted, and 39 completed the survey. Demographics of the total respondents are shown in Table 1; not all respondents completed every question.

### Survey Data analysis

All responses were pooled for initial data analysis. Descriptive and correlative statistics were generated for Likert-scale responses. There were three open-ended questions that included: “Why did you want to support a student or students through the publication process (writing, revision, and publication of the paper)?”, “What, if anything, did you learn about the following processes (research, writing, revision, and publication) during your mentoring of [Journal] students?”, and “What challenges did you face while guiding your student’s/students’ inquiry and writing?” There were 58-62 responses to each of the three open-ended responses on the survey. The open-ended responses were inductively coded in NVivo using constant comparative methods (Glaser, 1965). The developing codebook was discussed by members of the research team at weekly meetings, with subsequent revisions made and re-coding of responses as necessary. This process was repeated until we were unable to develop new codes and consensus was reached regarding the clarity of the codebook. The reliability of the coding framework was examined through an interrater reliability test of about 20% of the data with the first author. For codes that received a Cohen’s Kappa score below 0.3, the coders discussed the discrepancies, and the first author adjusted the codebook to reflect the consensus reached (McHugh, 2012). Across all codes, the Cohen’s Kappa averaged 0.47 with an average of 92% agreement across the coders.

#### Interview protocol development

The interview was developed by two coauthors in close collaboration using a constructivist approach by Charmaz (2003) which explored and defined processes. Questions were developed based on our research questions, discussed, and refined in consultation with other members of the research group. The researchers considered the validity and reliability of the list of questions through focused discussion between the team of researchers. This interview protocol contained six themes that touched on the mentors’ motivation, experience and self-efficacy, challenges, process, learning, and beliefs within the mentorship of science writing for young students.

### Interviews participants and setting

Interviews were conducted in this study following a survey to mentors of students who published a paper with [Journal]. The survey was sent to all mentors of students who published a paper, and at the end of the survey, the participants were asked if they were willing to participate in an interview, with a $20 Amazon gift card for their time. Of the 79 mentors who completed the survey, 14 mentors indicated their willingness to interview, and 13 were successfully scheduled (Table 2).

### Interview data collection and analysis

Semi-structured interviews were conducted by Zoom in Spring of 2021. Each interview was recorded over Zoom and transcribed, resulting in approximately 780 minutes of audio. The transcripts were deidentified. Mentors interviewed were asked if they had a preferred pseudonym, which we used, and others were assigned. Transcripts were initially read by [author 3] to develop emerging themes in Nvivo using open coding (Leech & Onwuegbuzie, 2011). Through further close reading of the text, [author 3] developed an initial set of codes. Coded text was compared, and their meanings discussed at weekly meetings by part of the research team. This involved thoroughly reading and comparing segments within and across categories to achieve consensus on interpreting the data and identifying emerging categories (Glaser, 1965). Following the initial codebook development[author 1] read through the transcripts and refined, added, and deleted codes. During weekly meetings the research team discussed the codebook and made revisions to eliminate inconsistencies or areas of confusion. This process occurred until the first author was unable to develop new codes. Codes were then grouped into larger themes that addressed each of the research questions. Author 2 analyzed a portion of the data to determine the interrater reliability. Cohen’s Kappa was run to determine the level of agreement between the two researchers; codes with a Cohen’s Kappa below 0.3 were discussed by the researchers and changes were made to the definition of the codes or how text was coded so that all codes had a Cohen’s Kappa of at least 0.3 or higher, with an average of 0.64, and a minimum of 85% inter-rater agreement, with an average of 91%. *IRB* The study was approved by the Emory University Institutional Review Board (STUDY00000797).

## Results

In this study, we conducted surveys and interviews with mentors to determine perceptions of publication within the scientific enterprise for pre-college students, the processes they used for mentoring students in writing and publication, and the experiences they had going through the process. While interview questions were designed to assess whether mentors considered the role of disciplinary literacy practices in scientific inquiry unprompted, the survey questions directly asked about their perceptions of these practices. For each research question we first describe the applicable survey data followed by the interview data to provide more insight into the findings (Bazeley, P. 2012).

### RQ1: What are the mentors’ perceptions of the disciplinary literacy practices of peer-review and publication?

Mentor perceptions and beliefs of a field can influence how they approach certain aspects of the scientific process as well as how they discuss or keep hidden those aspects with their mentee (Veal, et al., 2016). Thus, understanding how mentors perceive the process of publication could provide insight and connection to the approach they take in mentoring students through the process. We were interested in mentors’ general views on peer review and publication, such as how they might value it, and their experience and degree of confidence in the process.

From the survey, we investigated mentors’ general perceptions of peer-review and publication. We find that mentors value the scholarly publication and peer review process very highly, with near-total agreement with all statements, and always a majority selecting “Strongly Agree” for all statements (Table 3). Additionally, the majority of mentors report a high level of confidence in supporting students in different disciplinary literacy practices (Table 4).

**Table 3:** Mentor Perceptions of Peer Review and Scholarly Publication from Survey

| *Question and Response* | *Total* |
| --- | --- |
| **Peer review in the publication process improves the accuracy of the science presented** |  |
| Agree (*of which, Strongly Agree*) | 62 (38) |
| Neither Agree Nor Disagree | 2 |
| Disagree (*of which, Strongly Disagree*) | 0 |
| **Peer review in the publication process improves the communication of the science presented** |  |
| Agree (*of which, Strongly Agree*) | 61 (38) |
| Neither Agree Nor Disagree | 2 |
| Disagree (*of which, Strongly Disagree*) | 0 |
| **Publication is important because it helps a scientist share his or her science with a broader audience** |  |
| Agree (*of which, Strongly Agree*) | 63 (46) |
| Neither Agree Nor Disagree | 0 |
| Disagree (*of which, Strongly Disagree*) | 0 |
| **Publication is important because the science in a paper could be used by another scientist in his or her project** |  |
| Agree (*of which, Strongly Agree*) | 61 (46) |
| Neither Agree Nor Disagree | 2 |
| Disagree (*of which, Strongly Disagree*) | 1 (0) |
| **Peer review and publication are essential parts of doing science** |  |
| Agree (*of which, Strongly Agree*) | 63 (48) |
| Neither Agree Nor Disagree | 1 |
| Disagree (*of which, Strongly Disagree*) | 0 |

**Table 4:**
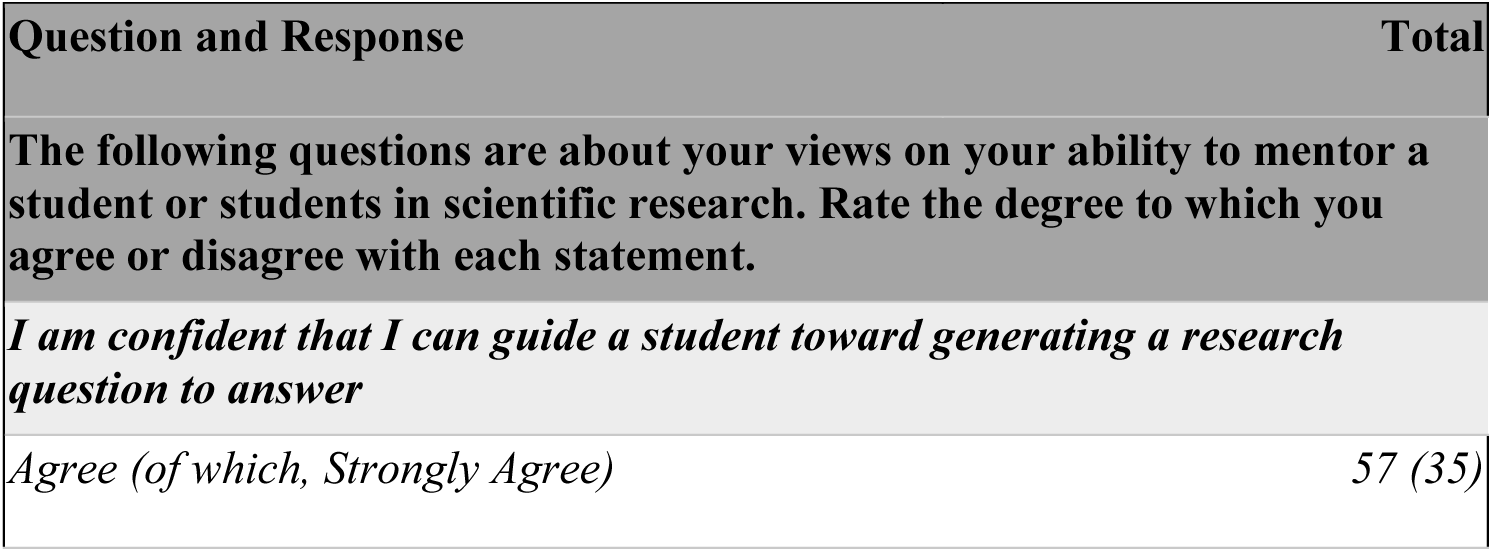
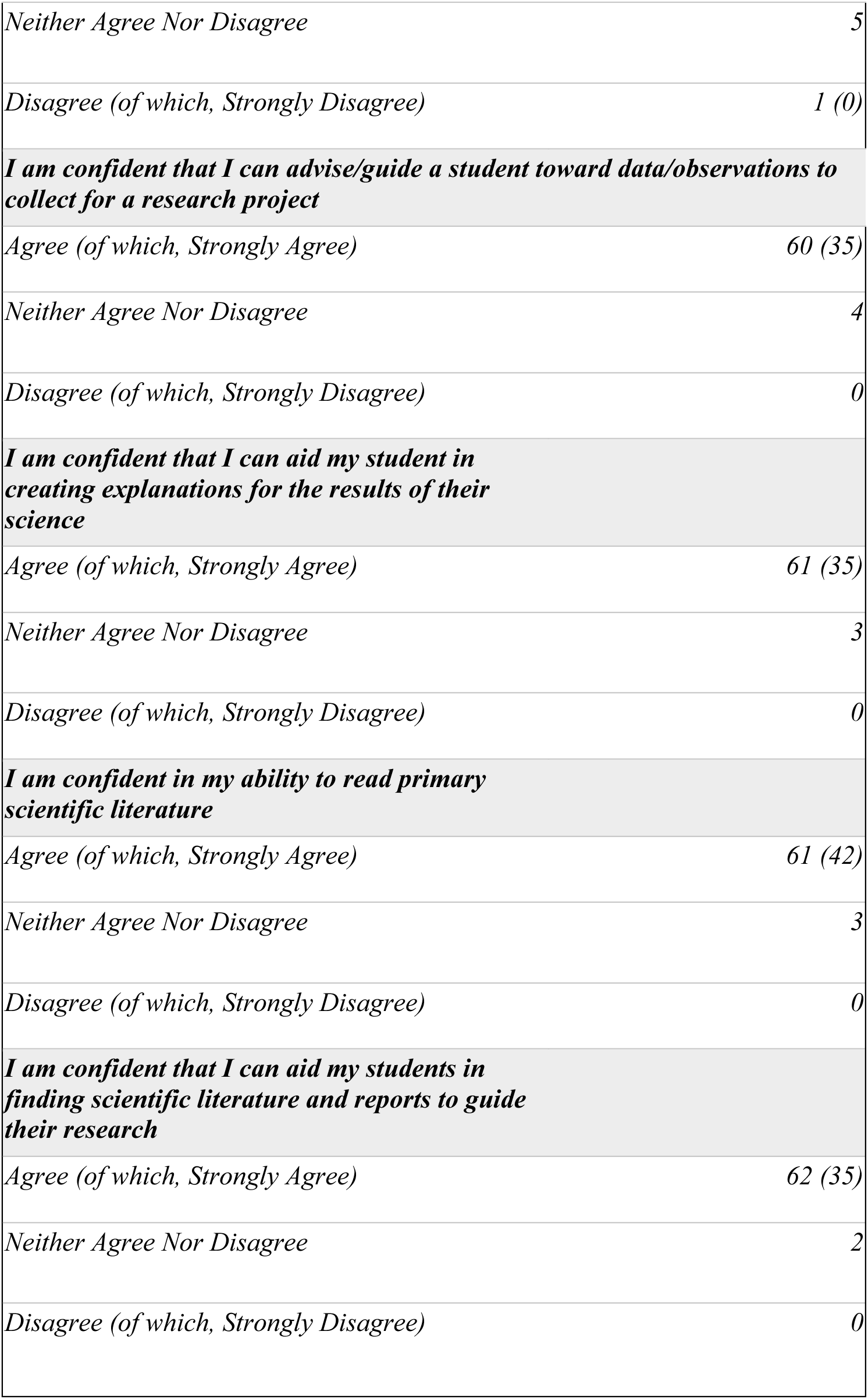
Mentor Self-reported Confidence in Disciplinary Literacy Practices

| Question and Response | Total |
| --- | --- |
| **The following questions are about your views on your ability to mentor a student or students in scientific research. Rate the degree to which you agree or disagree with each statement.** |  |
| ***I am confident that I can guide a student toward generating a research question to answer*** |  |
| Agree (*of which, Strongly Agree*) | 57 (35) |
| *Neither Agree Nor Disagree* | 5 |
| *Disagree (of which, Strongly Disagree)* | 1 (0) |
| ***I am confident that I can advise/guide a student toward data/observations to collect for a research project*** |  |
| *Agree (of which, Strongly Agree)* | 60 (35) |
| *Neither Agree Nor Disagree* | 4 |
| *Disagree (of which, Strongly Disagree)* | 0 |
| ***I am confident that I can aid my student in creating explanations for the results of their science*** |  |
| *Agree (of which, Strongly Agree)* | 61 (35) |
| *Neither Agree Nor Disagree* | 3 |
| *Disagree (of which, Strongly Disagree)* | 0 |
| ***I am confident in my ability to read primary scientific literature*** |  |
| *Agree (of which, Strongly Agree)* | 61 (42) |
| *Neither Agree Nor Disagree* | 3 |
| *Disagree (of which, Strongly Disagree)* | 0 |
| ***I am confident that I can aid my students in finding scientific literature and reports to guide their research*** |  |
| *Agree (of which, Strongly Agree)* | 62 (35) |
| *Neither Agree Nor Disagree* | 2 |
| *Disagree (of which, Strongly Disagree)* | 0 |

While the survey results suggest that mentors value, and are confident in, the publication process in general, the interview data provided more insight into how mentors conceive of the publication process within science, and in particular for students. The interviews were designed to investigate if communication- or publication-related responses would be elicited when mentors discussed their conceptions of doing science. For example, we asked mentors to “Explain your definition of science inquiry or the science process” and discuss what “skills are necessary to do science?” Of the 13 total mentors interviewed, six (5 TPMs and 1 SPM) had responses expressing their views on publication within the research process (Code: Views on Publication; Table 5). Five mentors discussed how publication plays an integral role in the expansion of scientific knowledge (Code: Expansion of Scientific Work; Table 5). For example, one mentor stated that “we want to share the idea to the world so that different scientists can know what they’re doing” (Clint). These mentors expressed the ideas that publication fosters collaboration and allows the scientific community to remain informed and interconnected. In addition to the expansion of scientific knowledge, three mentors were also found to express a “publish or perish” sentiment, where they acknowledged that the process of doing research needed to end in publication (Code: Publish or Perish: Table 5). These mentors espoused recognition for the notion that publishing is essential for survival in academia. As one TPMexplained: “Sadly enough, it’s like, you know, if you are a doctor, you need to see a patient otherwise, you’re not going to survive. And if you are a mechanic, you have to repair, if you don’t get any repair jobs, you’re not going to survive. That is the same case with this. If you publish, you are going to survive” (Bob).

**Table 5:** Mentor Perceptions of Peer Review and Scholarly Publication from Interviews

| Code | Definition | Total Number of Mentors out of 13 | Example |
| --- | --- | --- | --- |
| Expansion of Scientific Work | Mentors described publication as important because it involves the sharing of scientific work with other scientists around the world. | 5 | "The goal of science is to share the knowledge that you've come up with. So if you're not publishing it. How is anyone else going to know what you've learned?" (Hannah) |
| Publish or Perish | Mentors described publication as obligatory in the research process. They explain that all research must end in publication in order to be considered valuable. | 3 | "If you publish, you are going to survive, you are going to get more grants, you'll get more students and more of these that you can put all your questions in action. If you don't publish, you're not going to get any grants and then you may have really excellent questions but it will be just remaining as questions or somebody will steal it eventually" (Bob) |
| Quality of Publication | Mentors expressed that only high quality work should be published. They explain that only work that is significant and worth communication should be submitted for publication. | 2 | "I look at as the reviewer is whether it is a significant finding, whether it is worth publishing, whether it is worth communicating. If it is worth communicating, then I do all my help possible to get that paper published" (Bob) |

Two mentors also discussed the importance of publishing quality scientific work and ensuring that only accurate and significant findings are communicated (Code: Quality of Publication; Table 5). Overall, while the survey data demonstrates that mentors view the publication process as an important part of science, the interview data expands the understanding that mentors view publication as an important *output* of scientific research, more so than part of the process.

### RQ2: Why are mentors motivated to guide their students in publication?

Very little is known about *why* teachers or mentors would support high school student researchers through a voluntary and, at times, arduous process of publishing a research paper. From the survey, we analyzed responses from the open-ended question: “Why did you want to support a student or students through the publication process (writing, revision, and publication of the paper)?” Mentor responses were categorized into seven codes (Table 6). Analysis of the survey data’s most prevalent code revealed that 31% of mentors were motivated to guide their students through the publication process because they believed it was an important scientific experience for students to have (Code: Experience; Table 6). Many mentors with responses in this code focused on how engaging in peer-review and publication help students see the breadth of what science entails. For example, one mentor explained “I want students to understand the entire arc of the research experience from conceptualization to communication. I also want them to understand the intense and meaningful reality of what peer review really means.”

**Table 6.**
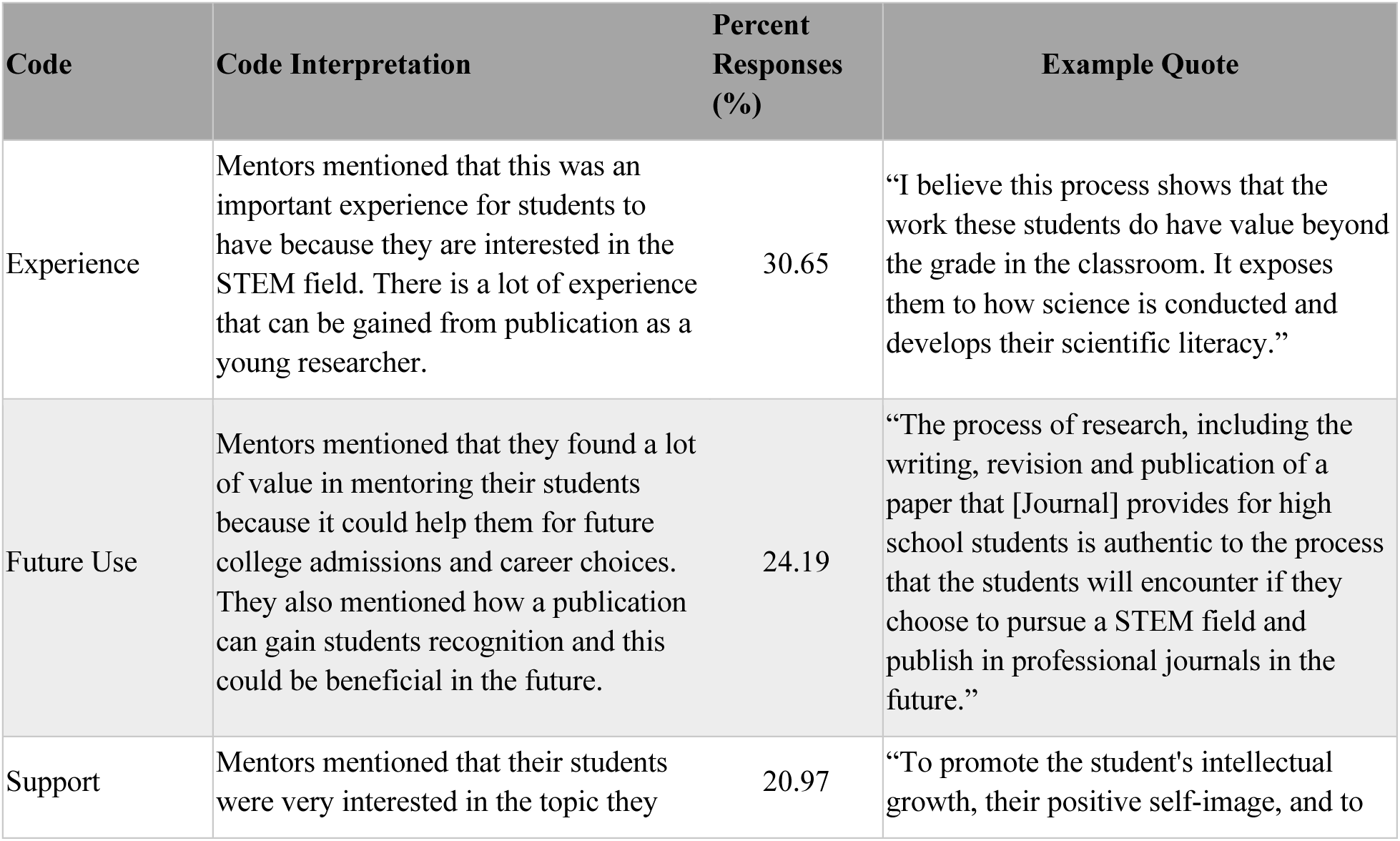
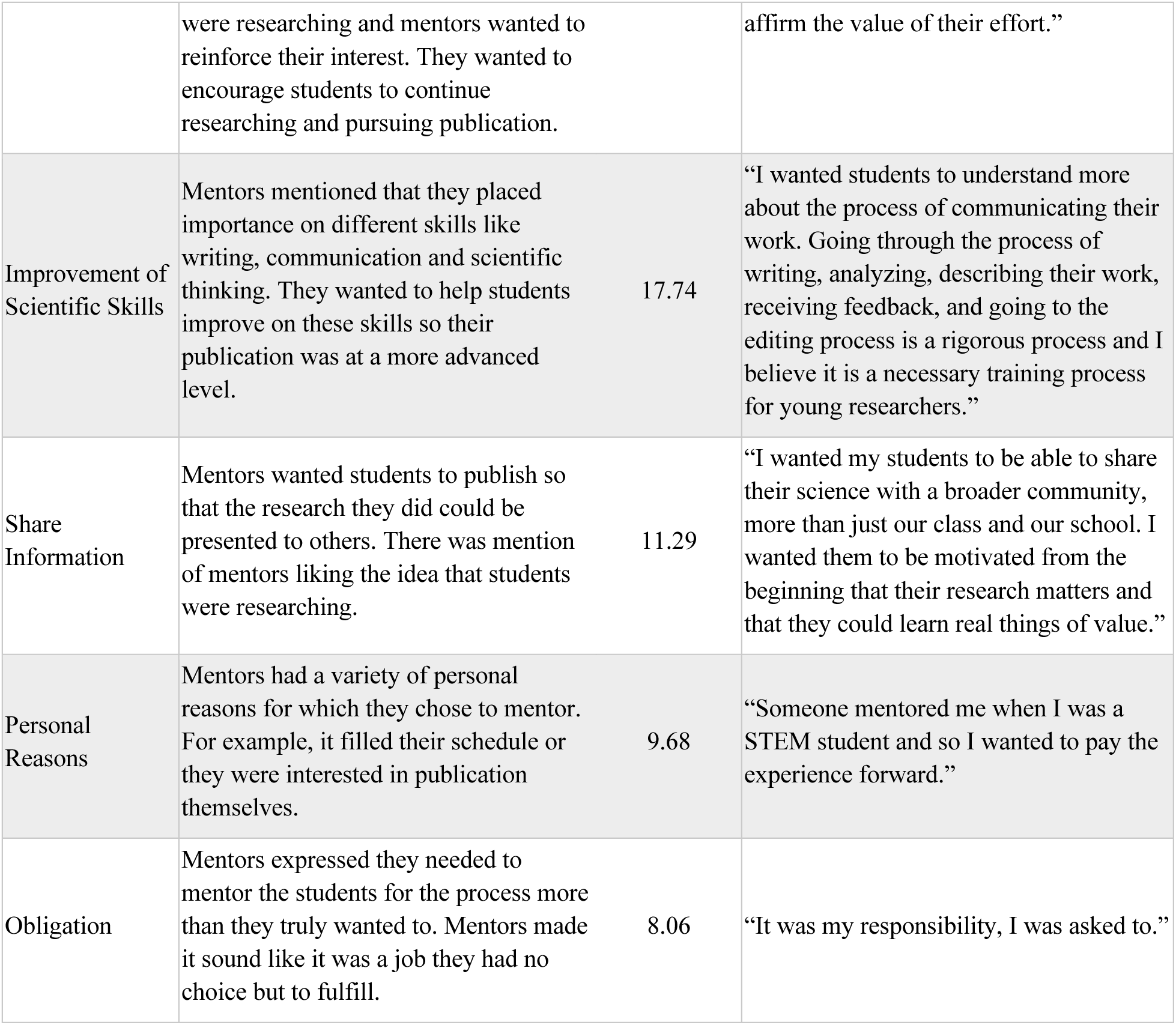
Survey responses to the question “Why did you want to support a student or students through the publication process (writing, revision and publication of the paper)?”

Additionally, 24% of mentors mentioned that they were motivated to mentor their students because it could help them with future college and career experiences (Code: Future Use; Table 6). For example, one mentor stated “…I believe that it prepares them for the rigors of college research and encourages them to remain in the field of research.” Besides college and career aspirations, but still under the topic of future use, mentors also suggested that a publication would help students gain recognition which would be beneficial for their future.

Interestingly, while the closed-response questions (Table 3) indicated that mentors believed the publication process is important for sharing work broadly, the open-ended responses revealed that only 11% of mentors expressed the idea that students should publish their work so that the research they did could be presented to others (Code: Share Information; Table 6). One mentor stated, “I wanted my students to be able to share their science with a broader community, more than just our class and our school.”

The interview data expands upon the themes revealed in the survey data (Table 7). Similar to the mentor responses in the “Experience” and “Future Use” codes of the survey data, the interview data demonstrated that a primary motivation for both TPMs and SPMs was to help students gain experience in peer review, with the majority of mentors, seven TPMs and four SPMs, mentioning this during their interview (Code: Experience in Peer Review; Table 7). These responses described how peer-review experience could provide students with valuable skills for the future and allow them to improve their scientific writing skills.For example, one mentor described, “I find that getting feedback from professionals is helpful for them, just getting them to be desensitized to it and not take it personally like, oh, this is a way I can make my writing better or more precise”(Hannah).

**Table 7.** Interview codes identified for mentor motivation for supporting students in publication process

| Code | Definition | Total Number of Mentors out of 13 | Example Quote |
| --- | --- | --- | --- |
| Experience in Peer Review | Mentors discussed how mentoring through [Journal] allows them to help students to gain experience in the peer review process. Experience in peer review was described as valuable due to its ability to motivate students, give students’ competitive advantage when applying to university, and enhance students’ skills in scientific writing. | 11 | "Oh, because I think it's like the pinnacle of doing research right. The pinnacle is to be published in a peer reviewed science journal. And the process of going through that peer review is super powerful and then having them actually see their work published. And then peer reviewed scientific journal is a huge deal. So I thought it was a great opportunity for my research students and you know they can it's like totally a feather in their cap" (Mallorie). |
| Engagement with Students | Mentors explained how connecting/supporting students was one of their reasons for becoming a [Journal] mentor. | 2 | "My main goal was just to support her, [student], and in whatever capacity, she wanted. I wasn't particularly pushing for to submit a big publication. I mean, I, the primary goal the research was just for this, the science fair that she did twice in two years in a row and So yeah I mean I viewed it you know once she decided she wanted to submit to this journal I viewed it as an opportunity for yeah, for me to understand the advisor side of someone's submitting the research project. And to, you know, give her experience with preparing the manuscript and going through the editing process, things like that" (David). |
| Personal Learning | Mentors described that they engaged in the mentorship process in order to gain mentorship experience and improve their scientific writing. | 1 | "Um well, for me personally, I definitely reading some of the papers and I was thinking about publishing, I just realized there was a lot I didn't know...So it was definitely there was a motivation for me too, because I was like, wow, like I think like if other high schoolers can do these things like with adults like I think my kids could do this and I think maybe I could so it definitely required I had to learn I had to learn a couple things first before I got confident enough to invite the kids to do it (Brian). |

Mentors not only explained their reasons for engaging in mentorship but also outlined specific goals that they wanted their students to achieve. Of the seven mentors who discussed outcomes for students, five TPMs discussed the goal of learning, and emphasized the importance of understanding peer review feedback (Code: Learning; Table 8). For instance, one TPM stated “I think trying to get them to really synthesize that feedback and then committing the time to really understanding it versus just doing what the teacher said” (Alex). Two TPMs also described competition as being one of their primary goals for students with one TPM stating “the goal is really for students to bring their research to science fair competitions”(Mallorie) (Code: Competition; Table 8). One SPM and one TPM described publication as their primary goal for students to instill a sense of seriousness and rigor in students. For example, the SPM stated “whatever you do it should be done, thinking that you will publish it one day…it has to be published because that will set a very serious tone to what you’re working on” (Shivani) (Code: Publication; Table 8).

**Table 8.** Interview codes identified for mentor goals for their students engaging in writing and publication of their scientific research papers.

| Code | Definition | Total Number of Mentors out of 13 | Example Quote |
| --- | --- | --- | --- |
| Learning | Mentors explained how they want students to learn from the [Journal] process. They explain how they want students to take ownership of the research process and synthesize the feedback they receive on their projects in order to maximize their learning. | 5 | "And I was like, look, you know, you might get published, you might not. If you don't, that's okay. That wasn't the goal of the experience. The goal of experience was for you to be able to take ownership of the whole arc of the research experience so that when you get into college. If you want to do this again, you're already building that skill set" (Hannah). |
| Competition | Mentors described how they aim to prepare students to present their research at science fairs and competitive platforms. | 2 | "And then, so the students always try to research plan, we always gear our research students towards ISEF, and so we use their paperwork and kind of get it in that direction" (Helen). |
| Publication | Mentors discussed how they want students to have an end goal of publication, and they explain how students should approach their research with a publication-oriented mindset. | 2 | "But if you are coming here, you need to go with that that mindset whether you publish it at the end or not is a different story, but at least you should have that mindset to communicate and that's how I started with it" (Bob). |

### RQ3: How do mentors support their students in peer-review and publication?

We were particularly interested in the ways in which mentors potentially enacted their views of writing and publication within science through mentoring their students through the process. Although the survey did not address details of the mentorship process, the interview provided details of the ways in which mentors supported students throughout the writing and publication processes. Mentor responses were analyzed and coded from the lens of cognitive apprenticeship. Three significant themes emerged which corresponded to the dimensions of “content”, “methods”, and “sociology”.

### How do mentors support their students in peer-review and publication: Content

The content dimension of cognitive apprenticeship refers to the distinct types of knowledge and thinking strategies that are needed for expertise. Mentors’ discussions revolved around two key principles within the "content’’ dimension of cognitive apprenticeship: “domain knowledge” and “learning strategies”. “Domain knowledge” pertains to subject matter-specific concepts, facts, and procedures, that students should know while "learning strategies" is about understanding how to help students grasp new concepts, facts, and procedures (Collins, et al., 1991; Minshew et al., 2021).

Mentors’ incorporation of “domain knowledge’’ was most evident when describing how they introduced students to scientific literature. Of the 13 total mentors interviewed, six TPMs and one SPM discussed “domain knowledge” by describing the strategies they used to help students gain proficiency in reading and finding scientific literature (Table 9). These mentors focused on the strategies of “finding literature” and “literature review” to guide students on how to approach scientific papers. All mentors who talked about teaching scientific literature as part of "domain knowledge" mentioned employing a “literature review” strategy as a mechanism to understand structure and central themes of scientific articles. For instance, one mentor explained that “ I select some uh papers on the [Journal] and print to them and I asked them to read it, and, for example, like, I gave them one paper per week and I want them to read it, and then, and then to tell me what is it about and uh at the very beginning it was very difficult and uh because there are lot of english that they don’t know and uh, for example, some chemical’s name. And they really don’t know unless they uh check the dictionary. But uh I- my opinion to them is just I just want them to um I don’t need to uh know what the scientific name is about, I just want them to see the structures of the paper” (Clint).

**Table 9.** Interview codes identified for mentor processes from the lens of cognitive apprenticeship

| Dimension of CA | Principles of CA | Total Number of Mentors out of 13 | Example Quote |
| --- | --- | --- | --- |
| Content | Learning Strategies | 7 | "...as they have questions. I often answer those questions with questions...So I often.. I'm pushing them to just continue to ask more questions and continue to figure out what resources they can use to answer those questions because that's what science is" (Mallorie) |
|  | Domain Knowledge | 7 | "I'll have kids go "There's nothing, I can't find an article about this." And I'm like, okay, let's sit down with our database. I want to see what search terms you used. I want to see like what Boolean operators, you used, right. I want to see what skills you have with keywords, etc. And then I'm like okay well I just found 2500 articles and you found zero. So we need to talk about how you're using that database" (Mallorie) |
|  | Control Strategies | N/A | N/A |
|  | Heuristic Strategies | N/A | N/A |
| Methods | Exploration | 13 | "Yeah I think they.. I think they will both learn a lot more and remember a lot more by going through it themselves and kind of leading.. It kind of helps you see the mentee see the big picture and not just okay, this is the task in front of me...I think having the mentee kind of find their own way, a little more helps them see the broader picture of okay, how does this minor task fit into my scientific study and how does the study fit into the field, and how does that fit into the world" (Brandon) |
|  | Coaching | 10 | "So usually what will happen and still have the kids work on it for like a week or so on their own and send it back to me. Then I'll give some comments back to them to sort of asynchronously we'll trade sort of ideas and feedback... Yeah. So I'm definitely involved, but I would say it's less hands on and sort of just like asynchronous feedback” (Brian). |
|  | Scaffolding | 8 | “So like before they want to write the manuscript I tell them the whole process of the publication thing, so I told him oh first you need to draft it and then you need to submit to the peer review and yeah I also tell them about the purpose of peer review, because we can’t just publish anything to the world and that we need to have some expert from that field who have some opinion on your project and the goal for this is to help you to improve and I told them that we definitely learned a lot from those peer review” (Clint). |
|  | Articulations | 5 | “they need to have a presentation skills so, for example, like uh I always ask my students to do presentations to me when um they finished doing the experiment, so uh they need to convey the idea to clearly to me” (Clint). |
|  | Reflection | N/A | N/A |
|  | Modeling | N/A | N/A |
| Sociology | Cooperation/Collaboration | 13 | “They do a fair amount of collaboration here and they will often many of their lab reports they actually write together as groups, so two or three kids will write it together and those groupings is collaboration sometimes go really well and sometimes you know terribly. And I think we talked, and then we'll talk about kind of what went well and what went terribly and then we also try to do a little bit of peer review in our classes where other groups will you know help other groups. So we try to kind of model some of that” (John). |
| Sequencing | Increasing Complexity | N/A | N/A |
|  | Increasing Diversity | N/A | N/A |
|  | Global to Local Skills | N/A | N/A |

Meanwhile, the strategy of “finding literature” was less prevalent, with only three TPMs discussing teaching students how to effectively search for primary literature using databases. There was a notable difference in the number of SPMs within the principle of “domain knowledge”. Only one SPM discussed the role of scientific literature in their students’ learning.

Within the context of “learning strategies”, seven TPMs described their methods for enhancing students’ reasoning abilities (Table 9), focusing on two key approaches: “feedback assimilation” and “inquiry”. Four TPMs employed the “feedback assimilation” strategy and explained how they encourage students to critically analyze and embrace the feedback they receive on their projects. One mentor explained, “So they’re taking what the edits are that are being given to them, and they feel like they have to just just follow what was given you know just change it, and I think that I want them more to think about why your suggestions are what they are, what they feel about it, and then to make the adjustment appropriately, you know, based on their opinions also” (Helen). Mentors also aimed to cultivate critical thinking among students through the “inquiry” approach where they motivated students to ask questions, for example one mentor mentioned: “We talked a lot about you know what type of study was…was their sample size, efficient, you know, what was the quality of their experimental design. Do you agree with their conclusions? How is this paper useful to helping you on your journey?” (Hannah).

### How do mentors support their students in peer-review and publication: Methods

The methods dimension of cognitive apprenticeship highlights the teaching strategies mentors employed to guide their students in the peer review process. All of the interviewed mentors, 8 TPMs and 5 SPMs, were found to use principles of “coaching”, “scaffolding”, “articulations”, “exploration”, and “modeling” within the methods dimension, with many mentors employing multiple methods.

Encouraging student independence was the most prevalent theme as is evidenced by the code “Exploration” (Table 9). These mentors emphasized the importance of intrinsic motivation and autonomy, maintaining lower levels of involvement with students in order to allow them to take initiative and ownership over their learning. One mentor described that “they’re driving the process… I’ll support them in any way but I pretty much hand it to them” (John). “Coaching” was found to be the next most prevalent principle employed, with ten mentors describing their approach of observing and facilitating while students carried out tasks. While they were invested in enhancing their students’ work, they also limited their involvement in order to preserve students’ independence and autonomy. Scaffolding was one of the least discussed principles among interviewed mentors as it was only expressed by eight mentors (Table 9). “Scaffolding" is when mentors offer structured support to help students perform tasks. These mentors described being more heavily involved in their students’ research endeavors, for example one TPM within this category explained that “I’m proofreading every section as they go along” (Hannah). Additionally, mentors’ involvement varied not just in principle but also in response to students’ individual capabilities and competencies. Three TPMs described how students’ writing abilities determined their level of engagement in the writing process; however, no SPM addressed using such adaptations.

Although less prominent, mentors also discussed how they encouraged students to communicate their research and scientific thinking as part of the research process. Within the context of “articulations”, five out of the eight TPMs explained the importance of students communicating their research effectively (Table 9). For instance, one TPM stated: “Get all the results and then communicate it properly. Communicate it effectively so that even your classmate can understand it at the same time as scientists can understand it as well… I let them go and compete with grad students in our department. If there is a poster competition, if there are no high school students, I just put them in and then they just go out and blow the field” (Bob).

### How do mentors support their students in peer-review and publication: Sociology

The sociology dimension of cognitive apprenticeship refers to the formation of a cooperative learning community. Within the mentor process, one of the most prevalent themes was the principle of “cooperation/collaboration”, which encompassed sentiments related to students working together, or with others, to accomplish their goals. All interviewed mentors highlighted the importance of collaboration, focusing on two key aspects: encouraging students to interact with other scientists and cultivating their own relationships with students (Table 9). Eleven mentors discussed how they encouraged students to engage with peers, researchers, and reviewers in order to foster a dynamic scientific learning experience.

For instance, one mentor stated that “I try to teach them early on to communicate with subject matter experts. Okay. Well, you’ve been reading enough in your lit review. There are probably some names that have come up a few times. Okay well email that person ask them a question…several of them have offered to do phone calls or Zoom’s or multiple sessions where they just talk through kind of research methods and ideas and so I think that piece of the experience has been really profoundly helpful for my kids to see that scientists are collaborative” (Hannah).

Furthermore, mentors also elaborated on the collaboration they themselves maintained with their students. Within the context of “cooperation/collaboration”, there were seven mentors that reflected upon their evolving relationship with students and discussed how they adopted a collaborative partnership with their students. The majority of these mentors discussed how they developed a relationship with students that fostered the development of their independence and self-efficacy. One mentor explained that: “I am no longer the person evaluating them and I am just someone to help, I’m just someone to support them. So, it’s turned into a very different and a very positive relationship in that there’s no grade… they essentially will bounce ideas off of me and I’ll kind of have a back and forth. And then I’ll say, but it’s ultimately up to you. And they’ll kind of they they’re driving the process” (John).

### RQ4: What do the mentors experience throughout the process in terms of their own development? What outcomes do they experience?

The findings related to this research question revealed much about the experience that mentors had with their students, as well as their own learning. The open-ended question from the survey asked mentors: “What, if anything, did you learn about the following processes (research, writing, revision, and publication) during your mentoring of [Journal] students?”. Analysis revealed that 45% of mentors learned the most about the general publication and peer review process (Code: Publication and Peer Review Process; Table 10). They mentioned that they were surprised from the level of difficulty they faced and from how time-consuming the process was. They also gained new experiences and realized the importance of peer review for students. One mentor mentioned “I learned most about the publication process through working with my student that published in [Journal]…”.

**Table 10.** Survey responses to the question "What, if anything, did you learn about the following processes (research, writing, revision, and publication) during your mentoring of [Journal] students?”

| Code | Definition | Percent Responses (%) | Example Quote |
| --- | --- | --- | --- |
| Publication Process and Peer Review | Mentors gained lots of knowledge about how the process of publication works. They mentioned that they were surprised from the level of difficulty they faced and from how time-consuming it was. They also gained a lot of experience and realized the importance of peer review for students. Majority of mentors mentioned that they were previously unfamiliar with the [Journal] platform and they learned a lot about how publication through [Journal] works. | 44.83 | “This gave me more experience with the publication process” |
| None | Mentors felt that this experience did not teach them any valuable information as many of them had already published in the past. | 32.76 | “None. I have published throughout my career.” |
| Writing | Many mentors gained more knowledge on scientific writing specific to publication which they realized was different than what they had past experience with. They also mentioned that there were new writing standards they were not familiar with. | 17.24 | “I learned how to teach my students to write the science research paper properly.” |
| Experimental Design | Mentors mentioned that they learned about aspects of the research process they were not familiar with before like statistical techniques, experimental controls, and scientific writing. | 15.52 | “The revision process pushed me to learn some new statistical techniques for analyzing data.” |
| Uncodable | These responses did not answer the question asked regarding what the mentors learned from the publication and peer review mentoring process. | 3.45 | “I have helped other students to create research papers and only the very best do I encourage help my students to contact [Journal].” |
| Research Topic | While mentors worked with their students, they learned a lot about the specific topic that the student was researching. They were not familiar with the scientific topics being investigated. | 1.72 | “...my students' research was outside my area of expertise so it was a very interesting experience to work on something that I am not familiar with...” |

The next most prevalent theme, which included 34% of responses, revealed that mentors believed they did not learn anything new from the experience (Code: None; Table 10). In many of these responses mentors explained that they had gone through the publication and peer review process in the past, so they did not believe that this process taught them anything that they did not already know. Other codes that emerged included Experimental Design, Research Topic, Writing, and Uncodable, however, at much lower prevalences (Table 10).

Although the survey indicated that many mentors did not express learning anything from the publication process, the interviews revealed that many mentors believed that they had learned throughout the process. There were five TPMs and two SPMs that described how [Journal] engagement refined their mentorship skills, teaching them how to best guide students through publication (Code: Teaching; Table 11). Having gone through the publication process with students, one mentor explained how [Journal] allowed him to better define his role as a mentor: ”I found that yeah when doing such investigative study for them, they really need to have a high autonomy they need to have a freedom to choose what they want to do so, then my role will be to answer their question” (Clint). There were six TPMs and two SPMs that discussed how submitting and writing a manuscript with students refined their research skills (Code: Publication; Table 11). These mentors explained how working with students and receiving peer review feedback improved how they approach scientific writing: “having to teach someone the process or kind of coach them through it made me think more about like why I write things certain ways or how I should you know, maybe how I should write” (Brandon).

**Table 11.** Interview codes identified for mentor learning

| Code | Definition | Total Number of Mentors out of 13 | Example Quote |
| --- | --- | --- | --- |
| Publication | Mentors described the enhancement of their research and scientific writing skills through the process of submitting a manuscript with students to [Journal]. They explain learning from peer review feedback, which helped them refine their scientific writing and deepen their understanding of the manuscript submission process. | 8 | "seeing all of the different steps of the publication, the scientific review, which was really scientifically reviewed, which was really cool and then you know, moving on to the second. edits and then, following that was copy editing it was it was nice to see all of these different groups of people coming together to to work on this and it was nice for students to see that as well. So yes, that was all very new to me, whole publication process, which now having seen it a couple of times running through it's very logical and it makes sense. It's certainly have expanded my horizons on that" (Helen). |
| Teaching | Mentors explained how involvement in [Journal] improved their teaching strategies and allowed them to enhance their ability to support and facilitate students' scientific endeavors. | 7 | "Beyond that, I think I probably became a better teacher because it gave me time to think about the clearest way to explain things, you know, explaining regression to a 10th grader makes you think about how you're explaining it to a doctoral student. And sometimes Maybe they're the same level of consideration, sometimes they're not. So really maybe think about how I approach explaining some of the parts of the process, whether it was actual research methods or the publication process, but also trying to understand what motivates students. What doesn't motivate students. So I think that was helpful to me" (Alex). |

A final open-ended question was asked regarding the process the mentors went through: “What challenges did you face while guiding your student’s/students’ inquiry and writing?” For survey data, mentor responses were categorized into 7 subcodes under the theme of “Challenges” (Table 12). Analysis of these codes revealed that 31% of mentors found the most difficulties in dealing with the level of knowledge that students had before beginning this process (Code: Developing Student’s Knowledge; Table 12). Mentors commented frequently that students were confused regarding a variety of scientific processes including how to effectively conduct research, work with statistical techniques, and write scientifically. As one mentor noted, students seemed to “lack a lot of basic knowledge to conduct research all on their own”.

**Table 12.** Survey responses to the question “What challenges did you face while guiding your student’s/students’ inquiry and writing?”

| Code | Code Interpretation | Percent Responses (%) | Example Quote |
| --- | --- | --- | --- |
| Developing Student's Knowledge | Mentors mentioned that they had trouble working with the amount of knowledge that their students previously had coming into this process. Students were confused about how to conduct research, work with statistical techniques, and write scientifically. | 31.03 | "Students may lack a lot of basic knowledge to conduct research all on their own." |
| Time | Mentors mentioned that there were lots of time constraints and the whole process was time-consuming. | 22.41 | “A semester is not an extensive period of time to design, carry out, and communicate scientific research. So turn-around time of waiting to connect with subject matter experts, running up against supply issues or restrictions, can really make the writing and revision process quite compressed.” |
| Accessibility of Resources | Mentors mentioned that they had trouble gaining access to equipment and scientific articles due to financial issues or restricted access. | 15.52 | “Availability of research paper on a similar topic specially on the plant science.” |
| Involvement | Mentors mentioned that they did not know how much to help their students to ensure that the students were experiencing the whole process but also getting the assistance they needed. Mentors were confused about how to balance the amount of guidance they gave students. | 12.07 | “It was challenging to find the right balance between guiding the student in the inquiry/writing process versus letting them find their own way.” |
| Accessibility of Students | Mentors mentioned that they had trouble effectively communicating with their students due to reasons like they had graduated out of the class. | 10.34 | “The biggest challenges to the publication process occurred because the revisions needed to occur after the class was over: sometimes students had graduated, sometimes they had just moved on to other classes.” |
| Revisions | Mentors were surprised by the peer review feedback they received from [Journal]. They mentioned that they received heavy revisions and were not impressed with the reviewers. | 10.34 | “Some students had a hard time revising their manuscripts. Some of the comments were too hard for high school students.” |
| [Journal] Problems | Mentors mentioned that they had problems with the [Journal] platform. There were confusions with the interface and format of submission. | 6.90 | “the user interface with the [Journal] website was perhaps the most frustrating part of the process. Submissions to [Journal], including revisions and the entire process was extremely frustrating and nerve-wracking.” |
| Lack of Familiarity | Mentors mentioned that they themselves were not aware of how the publication process works, thus, it was challenging for them to be good mentors as they were learning along with the student. | 1.72 | “It was the very first time for me also. It was a completely new learning process.” |

Another prominent theme that appeared was time constraints and the sentiment that the publication process was time-consuming (Code: Time; Table 12). One mentor mentioned, “… turn-around time of waiting to connect with subject matter experts, running up against supply issues or restrictions, can really make the writing and revision process quite compressed.”

Although mentors expressed challenges in the process, the interview data revealed the value that mentors placed on the outcomes of the [Journal] experience (Table 13). Three SPMs described the value of the process due to its ability to have realistic expectations that hold students to a higher caliber and foster the production of quality work (Code: Professional Expectations; Table 13). For example, a SPM explained “how much similar” [Journal] was to a former publication process and described to her students that “people here are not treating you like you are a different kind of scientist… this is exactly the process as it is and they are taking it very seriously” (Rosa). Furthermore, four out of five SPMs expressed that they found the [Journal] process to be valuable because it provides students with a credible platform to conduct and present their research (Code: Credible Platform and Recognition; Table 13). One SPM explained that “it provides a place for students to actually just you know do some do some sort of science that feels legitimate and real science like I remember” (Brandon). Compared to the SPMs, evaluation of the “value" code revealed that TPMs focused on the learning and engagement associated with peer review. When describing the most valuable aspects of the [Journal] process, seven TPMs and one SPM discussed how students could communicate with other scientists through participating in peer review (Code: Engagement with other Scientists; Table 13). One TPM explained how “I’ve commented on the kids papers and it’s always exciting when we get peer review comments back and someone suggests something or kind of asks a question where I’m just like, wow, like I did not think about that. So it’s been really helpful especially when it’s people who sometimes are a bit out of the field” (Brian). Overall, we find that SPMs tend to view publication through the [Journal] process as a means through which students can be held to higher scientific standards to produce credible scientific work, whereas the TPMs focus on learning through engagement with other scientists.

**Table 13.** Interview responses identified for mentor-identified value of the writing and publication process

| Code | Definition | Total Number of Mentors out of 13 | Example Quote |
| --- | --- | --- | --- |
| Engagement with other Scientist | Mentors discussed how participation in research through [Journal] facilitates students' engagement with the scientific community, allowing them to communicate with other scientists and receive feedback. | 8 | "I would say that just the process of going through collaboration with someone who is not your equal as opposed, you know, writing with your peers in high school is different than writing with someone who's either much more or even online, much less educated because you get an entirely different perspective. Sometimes it's those fresh eyes. They go, but wait..That's not right. Whether you have a PhD here or a 10th grade education or you're from here, you're from there, or your whatever your demographic characteristics are. There's always something new and what you know, use your intellect to so so fresh and different that participating, that process is just this beautiful thing of collaboration and learning from other people" (Alex). |
| Credible Platform and Recognition | Mentors explained how [Journal] is valuable in providing a credible and legitimate platform for students to conduct and present their scientific research. | 5 | "Yeah I mean like I said, I think, the one thing that really pleasantly surprised me about the process with [Journal] was the fact that it's a real journal and it's you know it's publishing real scientific data it's not just this little kids thing that you know nobody's ever going to read. It's real and the process with real and the reviewers were real and they they really they looked hard at what we've done and they made some excellent suggestion" (Adam). |
| Professional Expectations | Mentors explained that the [Journal] process holds students to a higher caliber, fostering the production of quality, professional-level scientific work that is feasible for a high school student to achieve. | 3 | "It has to be good, and it has to be done realistically, and that is why I think I like the system of [Journal]. And the fact that they encourage high school students to do and their expectations for a high school student is very realistic" (Shivani). |

## Discussion

Engaging the high school student in authentic science writing and publication, while relatively limited, is a growing experience. We anticipate that such student engagement not only serves to elevate their science literacy (Golden, 2023; Fankhauser, et al 2021; Mattison, et al 2022) but could also facilitate their development of identity and belonging in science (Carlone & Johnson, 2007; Deemer et al., 2022; Florence & Yore, 2004; Kim, et al 2023; Otto et al., 2023; Trujillo & Tanner, 2017). Thus, as these types of opportunities become more widespread, it’s important to understand how we’re mentoring our students to become familiar with the genre norms and the role of this type of communication within the process of science. Through surveys and in-depth interview data from mentors regarding their perceptions, we have developed several key conclusions:

### Mentors Express Multifaceted Perspectives on the Role of Publication

From the survey responses in RQ1, the vast majority of mentors emphasize the ideas that peer review and publication are essential parts of *doing* science and that publication is important for sharing work broadly with the community, with the majority of mentors saying that publication improves both the communication and the accuracy of the science presented. This signifies not only that mentors believe the research process is important, but also its effective communication with the scientific community as equally essential.

Somewhat in contrast, the interview responses revealed that mentors often view publication as a research outcome, with some acknowledging the prevailing "publish or perish" notion in academia and highlighting the necessity for research to culminate in publication. Combining insights from both the survey and interview data reveal that when mentors are presented the option that couches publication as part of the scientific process, the majority of mentors select the option. However, when unprompted, the mentors focus on more traditional views of publication as an output of research. It is likely that mentors’ views on publication are more nuanced and not limited to “publish or perish” sentiments. However, it raises questions of how mentors are discussing the role of writing and publication within science with their students.

When discussing science publication specifically for students, mentors almost exclusively focused on the value of the peer review process for their students, rather than mentioning other reasons for students to engage in publication (Tables 7 and 8). This focus likely helps contextualize the peer review practice for students, as long as mentors are articulating their value of the process for students.

However, unlike the cognitive apprenticeship model, which seeks to engage students in the authentic and professional practices of peer review and publication, mentors’ views and values about the publication process in RQ1 do not fully align with the goals they discussed for their students in RQ2. These siloed perceptions suggest that mentors may not be fully engaging students in what they perceive to be the professional practices and frameworks of thinking within the scientific profession, suggesting room for exploration on how to better immerse students in more authentic ways of thinking about the role of writing and publication within science.

### TPMs and SPMs Express Distinct Mentorship Goals

Furthermore, in RQ2, the interview data also highlighted the differences in goals mentors held for their students. Only one SPM explicitly discussed their student goals, emphasizing the importance of maintaining publication-oriented goals in order to cultivate a sense of seriousness and rigor. In contrast, the majority of TPMs discussed their goals for students. While TPMs were more vocal about the value of the publication process, SPMs were less so and tended to focus more on the outcomes of publication. This contrast potentially explains the observed discrepancy between mentors’ general views on publication and their goals for students as indicated in the survey data. Perhaps the fundamental goals driving mentorship approaches may vary between TPMs and other types of mentors. Previous studies have demonstrated that different forms of mentorship can profoundly affect the development of students’ scientific identity (Robnett et al., 2018). Furthermore, research by Fortus and Touitou (2021) demonstrates that the messages mentors relay to students, both implicitly and explicitly, impact students’ motivation in STEM. Thus, recognizing the distinct goals and motivations of mentors is crucial for a more accurate assessment of how mentors may influence students’ perceptions of the research process. As highlighted in our earlier research question, understanding the differences in mentors goals and perceptions can develop more cohesive and unified mentorship programs, ensuring consistent support for students across various mentoring relationships.

#### TPMs and SPMs have Divergent Mentorship Approaches

While all mentors were found to employ aspects of the cognitive apprenticeship model, the way in which they did so differed between TPMs and SPMs. The distinct backgrounds and environments of these mentors were found to largely shape their underlying values concerning the research process, which in turn, informed the specific goals and strategies they adopted while guiding students.

We found that the majority of TPMs interviewed engaged with students through school-based research curriculums (Table 2). Many of these TPMs reported that their students began the [Journal] program with a foundational understanding of the research process, setting the stage for advanced skill development. This contrasts with the survey data, where mentors expressed students’ limited research knowledge as a challenge (Table 12).This discrepancy may arise because the survey data does not distinguish between TPMs and SPMs, whereas discussions about students’ research experience and foundational knowledge were exclusive to TPMs in the interview data. Previous research has demonstrated how research curricula are designed to help prepare students for real-world scientific research and independent problem solving, with many scaffolding research skills to better equip students with the necessary knowledge to engage in independent research (Morrison et al., 2020). Therefore, TPMs working within such structured research curricula find themselves in an environment that is distinctly oriented towards student learning and are uniquely positioned to nurture essential skills in students. On the other hand, SPMs often engage with students independently, come from diverse professional backgrounds, and don’t have the framework of a formalized research curriculum (Table 2). As a result, they might not share the same educational emphasis as TPMs who operate within school-based research curricula.

RQ4 reveals how this divergence in the experiences and professional environments of mentors may shape their values regarding the research process and lead to distinct priorities in their mentorship approaches. TPMs, who might have less direct experience with the publication process, were not found to emphasize publication outcomes as heavily as SPMs. Instead, they prioritized the educational aspects of research, such as the pedagogical benefits of peer review and the learning that occurs through engagement with the scientific community. Conversely, SPMs placed a greater emphasis on the tangible outcomes of the research process, such as the recognition, platform, and quality assurance that publication provides for the students’ work. This difference in values not only reflects the mentors’ own professional experiences and environments, but also the goals they set for their students and their own personal reasons for becoming a mentor. As observed in RQ2, TPMs were more focused on learning-oriented objectives, while SPMs, though generally less explicit about their goals, had output-oriented goals.

These underlying values and goals likely also shaped the mentorship approaches used by TPMs and SPMs. In RQ3, within the “domain knowledge”, “learning strategies”, and “articulations” principles of cognitive apprenticeship, TPMs predominantly emphasized teaching students how to read scientific literature, develop critical thinking skills, and present their work. In contrast, only one SPM mentioned introducing students to reading scientific papers, and no SPMs discussed fostering critical thinking and presentation skills. This difference in approach reflects the unique sets of values held by mentors. TPMs primarily valued the learning-related aspects of the research process and in turn utilized principles of the cognitive apprenticeship model to cultivate essential skills among students. Thus, in RQ3, we see that differences in mentors’ values have tangible implications for students’ learning outcomes and their development through the mentorship process.

This context sets the stage for understanding mentors’ varied reflections on what they learned from engaging with [Journal]. In RQ4, all interviewed TPMs expressed what they learned from the research process, while only a minority of SPMs did the same. This discrepancy underscores how the values held by mentors not only shape their approaches to mentorship with students but also influence what they themselves gain from the research process. Furthermore, this observation potentially explains the results obtained from the survey where 33% of mentors revealed that they did not learn anything from the research process. Perhaps the extent to which mentors learn something from the [Journal] process depends on their professional background and values. SPMs may simply not value the pedagogical aspects as much as TPMs, and thus may be less likely to discuss how engagement in [Journal] influenced their own mentorship abilities.

Overall, throughout this study the differences in backgrounds between TPMs and SPMs emerge as pivotal factors shaping their distinct values and perspectives on the publication process. TPMs’ focus on learning and cultivating essential skills among students predominantly aligns with the "performance" and "competence" aspects of STEM identity, facilitating students’ engagement in essential scientific practices and enhancing their proficiency in these areas. However, SPMs’ emphasis on publication-related outcomes of the research process highlights the "recognition" aspect of STEM identity, where students’ participation in publication and peer review processes enables them to gain recognition for their work (Carlone and Johnson, 2007). As students navigate these different mentorship styles they can be influenced by the underlying values and perceptions embedded in these approaches. This exposure to different mentorship styles is a crucial part of the hidden curriculum and can lead to varying perceptions among students about what constitutes successful research (Orón et al., 2018). Moreover, this variation in perceptions raises important questions about the balance of these approaches in scientific education. It suggests a need for an integrated model that merges the strengths of both mentorship styles. Such a model would offer students a well-rounded and nuanced grasp of the research process while nurturing all facets of their STEM identity.

#### Mentors aim to Balance Autonomy and Guidance in Research Mentorship

Across the dimensions of "methods" and "sociology," a prevailing theme observed was the encouragement of autonomy and self-sufficiency by mentors. In the “methods” dimension, the majority of mentors expressed that they take on a lower degree of involvement to foster self-efficacy among students, further reinforcing this stance in “sociology” by describing how they cultivated relationships with students that bolstered independence. Overall, the majority of mentors interviewed adopted a "learning by doing" approach, as described by Chong & Mason (2021), McDowell et al. (2019), Gewin (2019), and van der Loo et al. (2019), where students are encouraged to take ownership of the writing and peer review process. Studies have indicated that promoting self-efficacy among students may increase their motivations to continue to pursue STEM career paths and may help them gain confidence in their research capabilities (Butz et al., 2018). Therefore, encouraging students to take the lead in their learning and research efforts may more effectively aid in the development of their STEM identity through fostering confidence and a lasting interest in academic research.

However, while fostering independence and self-efficacy among students may have positive impacts on students’ growth and science identity, this approach raises questions on equity and the expectation for students to be self-directed, especially in a process that they may be unfamiliar with. Not all students may have the prior experience and foundational skills necessary to independently navigate the research process. The survey data highlighted that one of the main challenges mentors faced was accommodating students’ confusion over the fundamental aspects of the research process. Teaching foundational skills seemed to be a significant part of the mentoring process demonstrating the importance of mentors filling in knowledge gaps and providing individualized support. Three of the interviewed TPMs expanded upon this notion, noting that they often needed to adjust their levels of involvement to better align with each student’s unique needs and capabilities. Furthermore, previous studies, such as Moreu et al. (2021), have underscored the significance of inclusive academic environments that cater to the varied needs of a diverse student body. Research by Ghazzawi et al. (2021) discusses how students from underrepresented groups often encounter unique challenges in STEM fields, which can impact their self-confidence and persistence in these disciplines. Likewise, research by Roehrig et al. (2021) advocates for an approach that directly guides students through authentic scientific practices as found in real-world scenarios, suggesting that this method can foster the development of positive STEM identities among underrepresented students. Therefore, while a "learning by doing" approach may offer several benefits, it’s essential to pair it with personalized mentorship that considers students’ unique needs and more explicitly highlights aspects of the research process that are part of the hidden curriculum. It’s important to note that despite the overarching theme of fostering independence among students, a subset of mentors did discuss delivering direct instruction to their students. TPMs expressed tailoring their levels of involvement based on student needs, and a subset of mentors employed the “scaffolding” principle within the “methods” dimension where they explained research concepts to students.

### Absence of the Sequencing Dimension of Cognitive Apprenticeship

Connected to the conclusions above regarding the balance between autonomy and guidance, mentors’ discussions did not encompass any aspects of the “sequencing” dimension of cognitive apprenticeship. The “sequencing” dimension revolves around the deliberate arrangement of activities from simpler tasks to the gradual increase in complexity to facilitate the development of expertise (Collins and Kapur, 2014; Minshew et al., 2021). For instance, with regards to scientific writing, sequencing could look like mentors guiding students to first outline their research papers, and then have them systematically focus on each individual section. In such an approach, providing students with a clear and logical progression of learning activities, may enhance students’ ability to connect different components of their projects and result in a more holistic understanding of the research process. It is important to note that the review conducted by Minshew et al. (2021), also observed that the dimension of “sequencing” tends to be the least discussed aspect within cognitive apprenticeship. This recurring observation suggests that aspects related to the “sequencing” dimension may generally be underrepresented in mentorship. This highlights the need for a refined approach to mentorship that better incorporates methods aimed at providing students with a well-structured framework for comprehending various aspects of their projects and the peer review process.

#### Limitations

While our results highlight the potential influence of current mentorship practices on students’ scientific identity, this study also has several limitations that underscore the need for further, more detailed exploration. A primary limitation is the small and uneven sample size of interviewed mentors. We included responses from only thirteen mentors, comprising eight TPMs and five SPMs. This discrepancy, along with the overall small sample size, may have skewed our analysis, particularly when comparing the mentorship practices between TPMs and SPMs. Additionally, there is a possibility that some mentors held specific beliefs or engaged in certain mentorship practices that they did not fully disclose during the interviews. This could potentially explain why there was little discussion about the sequencing dimension among mentors. It’s possible that mentors did incorporate aspects of the sequencing dimension in their approach while mentoring; however, these elements may not have been captured in our analysis due to the small sample size and limited scope of the surveys and interviews. Furthermore, given the moderate interrater agreement in the survey data, it is important to acknowledge that the qualitative analysis of open-ended survey and interview responses leaves room for interpretation, highlighting the potential variability in how mentorship practices are perceived and reported. Overall, these limitations suggest a need for broader studies to more accurately understand the range of mentorship objectives across different mentor types. Such studies would aim to expand the sample size and diversity of mentors, as well as employ methodologies that encourage more transparent and detailed sharing of mentorship practices. This could involve designing longitudinal studies where mentors are interviewed and observed for extended periods, providing more insight into the growth of their mentorship styles and the nuances of their approach.

### Conclusion and Future Directions

Our study suggests that mentors’ backgrounds and perceptions of publication influence their goals for students and shape their approach to mentorship. Our findings indicate that variations in mentors’ values and methodologies could yield diverse outcomes for students, underscoring distinct facets of their science identity. Mentors’ diverse views on the role of publication in research—from the experiential process to the value of the final outcomes—suggest the potential to nurture various aspects of students’ scientific identity, including their sense of performance, competence, and recognition in the field.

Likewise, our findings revealed distinct mentorship values and approaches between TPMs and SPMs. TPM’s discussions placed a larger emphasis on learning and skill development, which may showcase a desire to nurture the performance and competence aspects of students’ science identity. Meanwhile, SPMs’ interview responses focused more on publication outcomes, likely fostering the recognition dimension of a student’s science identity.

Furthermore, in our findings, we also identified areas for improvement in existing mentorship practices. Our analysis revealed discrepancies between mentors’ views on publication and their goals for students, suggesting a lack of immersion in ways of thinking in the discipline. To improve this approach, we recommend that mentors reflect on the goals they set for students and assess whether these align with their own views of authentic scientific practices. Additionally, while most mentors discussed fostering self-efficacy and autonomy among students, a lack of structured guidance, particularly in the research and publication processes, could hinder students’ integration into the scientific community (Moreu et al., 2021). This is compounded by the minimal focus on the "sequencing" dimension within mentorship, further suggesting that students may not be receiving structured guidance and support (Collins and Kapur, 2014; Minshew et al., 2021). Thus, we call for a more dynamic mentorship model in which mentors tailor the level of encouraged independence to each student’s background. Incorporating elements of the "sequencing" dimension, among others, provides students with more structured and direct support for scientific reading and writing which can enhance the performance and sequencing aspects of their science identity. This approach is especially more equitable for students from underrepresented backgrounds who may have had limited exposure to STEM fields and thus require more direct and systematic support.

Overall, adopting this tailored and structured approach to mentorship has the potential to integrate students into authentic scientific practices in a more equitable and inclusive manner.

Disciplinary literacy practices are a critical part of scientific research, and further research is needed for a more comprehensive understanding of the impact that mentorship approaches have on student outcomes in science disciplinary literacy practices. Future studies should aim to explore how mentorship practices influence students’ implicit beliefs and attitudes towards writing and publication, Examining student outcomes may facilitate a more precise evaluation of how current mentorship practices affect students’ STEM identity and their continued engagement in scientific fields (Atkins et al., 2020; Chang et al., 2011; Chemers et al., 2011; Hazari et al., 2013; Perez et al., 2014; Woods et al., 2023). This, in turn, may help identify potential areas for enhancing the efficacy of mentorship models.

## Supporting information

Interview Protocol

## Acknowledgements

We are grateful to Paige Crowl for her assistance in finding relevant literature for the paper. Additionally, we are grateful for the work of Devin Gee who provided feedback on the interview development and analysis.

