## Supplementary material for "Empowering future scientists: Mentors employ various strategies to engage students in professional science disciplinary literacy practices": Interview Protocol

Thank you for agreeing to be part of the mentor interviews that we are conducting. [introductions] First,  We will be using a recording device and the recording function of Zoom in order to make sure we do not lose any data. The recordings and all other data will be securely stored and only members of the research team will have access to them. Do you agree to being recorded via Zoom, recording device, or both? If you disconnect from the meeting please log back on using the same link.

We're conducting these interviews to get a sense of your experiences and views of science while mentoring a student through their *JEI* project. These interviews will supplement the data that we obtain from the mentor survey which has been sent to all *JEI* mentors, and we anticipate pursuing publication with the data we obtain. We do not expect any harm to come to you through these interviews. We have a few general questions and then follow up questions to ask you. The first set is about your background, the second set is about your experiences in mentoring a student through *JEI*, and the last set is about what you learned in the process. Your information will be kept confidential. Your data will be anonymized, and your research records will be purely confidential. So if you agree to do this study then may I get a verbal agreement from you now.

Do you have any questions for us before we begin? Do you agree to participate in this interview? **Tell us a little bit about your experience: how many students have you mentored through the *JEI* publication process?**

**1. What is the motivation for the research mentors in having students participate in the peer-review and publication process?**

- Why did you decide to mentor a student through the peer-review and publication process, and what, if any, goals did you have for yourself and/or your student for participating in the publication process? (is it required, checking a box, or something else?)
    - Who initiated the idea of publication with JEI ? Who decided on JEI  vs something else? (for all mentors)
        - At what point did you and your student decide to pursue publication of the project? (both)

- Did you feel comfortable mentoring your student's project, and was it in your certification subject? (more teacher)
  - Has your comfort level changed since mentoring the student or since going through the JEI process? (more teacher)

**2. (Confidence and self-efficacy and connection to teaching/learning) How much experience does the research mentor have with science writing (does that experience influence their learning, understanding, or teaching/mentoring of primary literature)? ( minutes)**

- How would you describe your familiarity with scientific writing and primary scientific literature? How did you gain this experience? (more teacher)
  - *Have you ever written a primary paper before?*
  - *Did this (familiarity/comfort/perception/knowledge) change during or after the project?*
  - *Do you think that mentoring a student helped you refine the skills of writing a primary article?*
- What is your level of confidence in helping a student who has little experience in reading and understanding primary literature in STEM? Did this confidence change during or after partaking in the project? (both)
  - *Do you feel more confident to help a student in the future through the JEI process?*
  - *Has this changed the way you mentor a student through research?*

3. **What challenges did the research mentor face in guiding student inquiry and/or writing? (minutes)**

- Overcome or not overcome, what did you find difficult about mentoring a student through the research or *JEI* publication process? (both)
  - *Which was the greatest challenge with mentoring a student through this process and why?*
  - ***(Maybe)****: How did you overcome these challenges?* or *Why were you not able to overcome them?*

4. **How much guidance did the mentor give the student in the writing process?( minutes)**

- How involved are you in the publication (writing, revision) process? (both)

- What is your level of confidence in helping a student who has little experience in reading and understanding primary literature in STEM? Did this confidence change during or after partaking in the project?
- Did you and your student discuss peer-review and publication as a part of doing science before or while working on the project? (both)
    - *What was your explanation of the role that peer-review and publication has in doing science?*
    - *If you were to explain these roles again with another student would this explanation change?*

5. **What did the mentor learn about the review and publication processes? (Optional question depending on answers in #4)( minutes)**

- After mentoring a student participating in *JEI* , how has your understanding of peer review and the publication process changed?
    - Community: Has your conception of how scientists can collaborate and cooperate changed at all?
    - What are unexpected outcomes of mentoring a student through JEI ?

6. **Has engaging in this process impacted their views and practice of scientific inquiry?( minutes) (BELIEFS—connected to practice/teaching)**

- What is your definition of scientific inquiry?
    - Did this definition change between the beginning of the project to the end of the project? (all mentors...it would be interesting to see if there is a difference)
- What are the characteristics of scientific inquiry?
- What kinds of skills are necessary to do good scientific inquiry?
- Why is scientific inquiry valuable? Who is capable of scientific inquiry?
    - Do you think this process (research, writing, publication) would be attainable or most of your students if they had the material resources? Do you think there is a certain "type" of student that *JEI* is for?
- How do the STEM disciplinary skills (reading, writing, publishing) fit into your view and practice of scientific inquiry, and has this changed after participating in JEI ?
- Would you like to add to or make changes to your definition of inquiry?
    - Have your ideas or thoughts on cooperativity and collaboration within science inquiry changed at all by going through the JEI process? (both)
- Did this mentoring experience affect your teaching or mentoring practice? (both)

   - How did this change your relationship with your student vs. another student who didn't go through JEI  but still participated in research? (optional if we have time)
   - What is one new thing you learned about the scientific process or scientific inquiry? (Test Question)

**7. Additional Questions( minutes):**

- What do you see as the value of JEI  or participating in JEI ?
- Would it be plausible and would you consider incorporating parts of the *JEI*  process into your curriculum? Why or why not? (teacher)

Please provide any additional comments on your experience with *JEI* .
